# Structural basis of human LRRK2 membrane recruitment and activation

**DOI:** 10.1101/2022.04.26.489605

**Authors:** Hanwen Zhu, Francesca Tonelli, Dario R. Alessi, Ji Sun

## Abstract

Mutations in *LRRK2* are the most common genetic cause of late-onset Parkinson’s disease (PD). *LRRK2* encodes the leucine-rich repeat kinase 2 (LRRK2), whose kinase activity is regulated by Rab29, a membrane-anchored GTPase. However, molecular mechanisms underlying Rab29-dependent recruitment and activation of LRRK2 remain unclear. Here we report cryo-EM structures of LRRK2–Rab29 complexes in three oligomeric states, illustrating snapshots of key steps during LRRK2 membrane recruitment and activation. Rab29 binds to the ARM domain of LRRK2, and disruption at the interface abrogates LRRK2 kinase activity. Activation of LRRK2 is underpinned by the formation of an unexpected Rab29-induced super-assembly containing two central kinase-active and two peripheral kinase-inactive LRRK2 protomers. Central protomers undergo pronounced oligomerization-associated rearrangements and adopt an active conformation. Our work reveals the structural mechanism for LRRK2’s spatial regulation controlled by Rab GTPases, provides mechanistic insights into pathogenic mutations and identifies new opportunities to design LRRK2 inhibitors for PD treatment.

## Introduction

Parkinson’s disease (PD) is the second most prevalent neurodegenerative disorder, affecting 1-2% of the population above the age of 65 (Lees et al., 2009). Mutations in the *LRRK2* gene, which encodes the leucine-rich repeat kinase 2 (LRRK2) protein, are one of the most frequent genetic causes of PD and account for ∼5% of familial and ∼1% of sporadic cases (Paisan-Ruiz et al., 2004; Tolosa et al., 2020; Zimprich et al., 2004). More than 250 mutations in *LRRK2* have been identified, but pathogenicity has been confirmed only for a small subset, including N1437H, R1441C/G/H, Y1699C, G2019S and I2020T (Blanca Ramirez et al., 2017; Bryant et al., 2021; Kalogeropulou et al., 2022). LRRK2^G2019S^, the most common mutation in LRRK2-related PD cases, significantly increased the kinase activity (Jaleel et al., 2007). Therefore, LRRK2 kinase inhibitors are of great interest in PD treatment (Tolosa et al., 2020). Understanding how LRRK2 functions and what goes wrong in disease-causing mutations will inform the rational design of specific LRRK2 inhibitors. Indeed, much effort has been directed at exploring the structure-function relationships in LRRK2 to gain a mechanistic understanding and guide the rational drug discovery (Usmani et al., 2021).

LRRK2 is a 286-kDa protein with seven domains, including armadillo repeats motifs (ARM), ankyrin repeats (ANK), leucine-rich repeats (LRR), Ras of complex (ROC), C-terminal of ROC (COR), kinase (KIN) and WD40 domains. We recently reported high-resolution cryo-EM structures of full-length human LRRK2 (Myasnikov et al., 2021), revealing the overall architecture. Like catalytic LRRK2^RCKW^ (Deniston et al., 2020), the full-length LRRK2 was also captured in an inactive state, whose features indicated LRRK2 activation would require considerable structural rearrangements. An active conformation has been observed by the *in-situ* cryo-ET analysis of a microtubule-associated filament assembly formed by the pathogenic LRRK2^I2020T^ (Watanabe et al., 2020). However, the 14-Å resolution precluded detailed insights into LRRK2 kinase activation.

Moreover, LRRK2 activation under physiological conditions is regulated through membrane recruitment and higher-order oligomerization, the structural basis of which remains yet unknown.

Rab GTPases are master regulators of intracellular vesicle trafficking and play a crucial role in the LRRK2 signaling as both upstream regulators and downstream substrates (Berwick et al., 2019). Rab29, encoded in the PD locus, *PARK16* (Tucci et al., 2010), belongs to the Rab GTPase family, a subgroup of which are well-characterized LRRK2 substrates including Rab8A, Rab10 and Rab29 (Steger et al., 2017; Steger et al., 2016). Prenylated Rab29 is localized at the Golgi apparatus and controls the membrane recruitment of LRRK2 in a GTP-dependent manner (Kuwahara and Iwatsubo, 2020; Liu et al., 2018; Purlyte et al., 2019). Rab29 recruitment stimulates the kinase activity of LRRK2, leading to increased phosphorylation of Rab29 and Rab10 and LRRK2 autophosphorylation at Ser1292 (Purlyte et al., 2019). In addition, directing Rab29 to other cellular membranes, including mitochondria, also induces LRRK2 recruitment and activation (Gomez et al., 2019). These data suggest an interesting spatial regulation mechanism of LRRK2, where Rab29-dependent membrane recruitment boosts LRRK2 kinase activity.

The kinase activity of LRRK2 could also be modulated by protein oligomerization (Civiero et al., 2017; Gloeckner et al., 2006; Greggio et al., 2008; Jorgensen et al., 2009). Previous studies indicated that LRRK2 could form homodimers via its COR or WD40 domains (Berger et al., 2010; Civiero et al., 2012; Deniston et al., 2020; Greggio et al., 2008; James et al., 2012; Klein et al., 2009; Sen et al., 2009; Watanabe et al., 2020). Moreover, higher oligomers of LRRK2 proposed to be critical for its kinase activity has also been reported both *in vitro* and *in vivo* (Berger et al., 2010; Greggio et al., 2008; James et al., 2012; Sen et al., 2009), but the molecular relationship between LRRK2 activity and protein oligomerization under normal physiological conditions remains puzzling. Here we determined the cryo-EM structures of LRRK2–Rab29 complexes in three different oligomerization states and, combined with structure-guided functional analyses, revealed unprecedented mechanistic insights into LRRK2 membrane recruitment and activation.

## Results

### Structural basis of Rab29-dependent recruitment of LRRK2

We first examined the interaction between GTP-bound Rab29 and full-length LRRK2 using an *in-vitro* GST pull-down assay and verified that wild-type Rab29 robustly interacted with recombinant LRRK2 (Figure S3E) (McGrath et al., 2019). Next, we reconstituted the complex by mixing LRRK2, Rab29, non-hydrolyzable GTP analog (GppNHp), ATP and Mg^2+^, and performed single-particle cryo-EM studies. Three-point mutations were introduced into Rab29 to prevent its phosphorylation by LRRK2 (T71A and S72A) and stabilize a GTP-bound conformation (Q67L), thus increasing the homogeneity of LRRK2–Rab29 complexes. The final cryo-EM reconstruction resulted in LRRK2– Rab29 complex structures in three distinct oligomerization states (Figures S1 and S2). Oligomerization state 1 (LRRK2 monomer state) contains one LRRK2 and one Rab29 (Figure S1B); oligomerization state 2 (LRRK2 dimer state) has two LRRK2 and two Rab29 (Figure S1B), and oligomerization state 3 (LRRK2 tetramer state) shows visible cryo-EM densities for four LRRK2 and two Rab29 molecules (Figure S1B). Focused 3D refinement was performed to improve the cryo-EM density surrounding the LRRK2–Rab29 interface (Figure S1A), and C2 symmetry was imposed for oligomerization states 2 and 3 during data analysis (Figure S1A and Table S1).

We will use the LRRK2 monomer state to dissect Rab29-dependent recruitment. LRRK2 in the oligomerization state 1 is almost identical to the inactive LRRK2-alone structure (Figure S4C) (Myasnikov et al., 2021). Rab29 binds to the N-terminal ARM domain of LRRK2, burying a surface area of ∼800 Å^2^ (Figure 1A), and adopts a “Switch I” closed configuration, which is usually shared by GTP-bound small GTPases (Figures S3A and S3C). No apparent nucleotide density was observed within Rab29 despite the presence of 0.5 mM GppNHp (Figure S3A), which may be due to a low binding affinity between this GTP analog and Rab29. Nevertheless, docking of the GDP-bound Rab29 structure (PDB 6HH2) (McGrath et al., 2019) into the cryo-EM density shows obvious steric clashes with LRRK2 (Figure S3D), rationalizing why the GTP-bound state promotes the Rab29-dependent recruitment of LRRK2 (Gomez et al., 2019; Liu et al., 2018; MacLeod et al., 2013; McGrath et al., 2019; Purlyte et al., 2018; Waschbusch et al., 2014).

**Figure 1.**
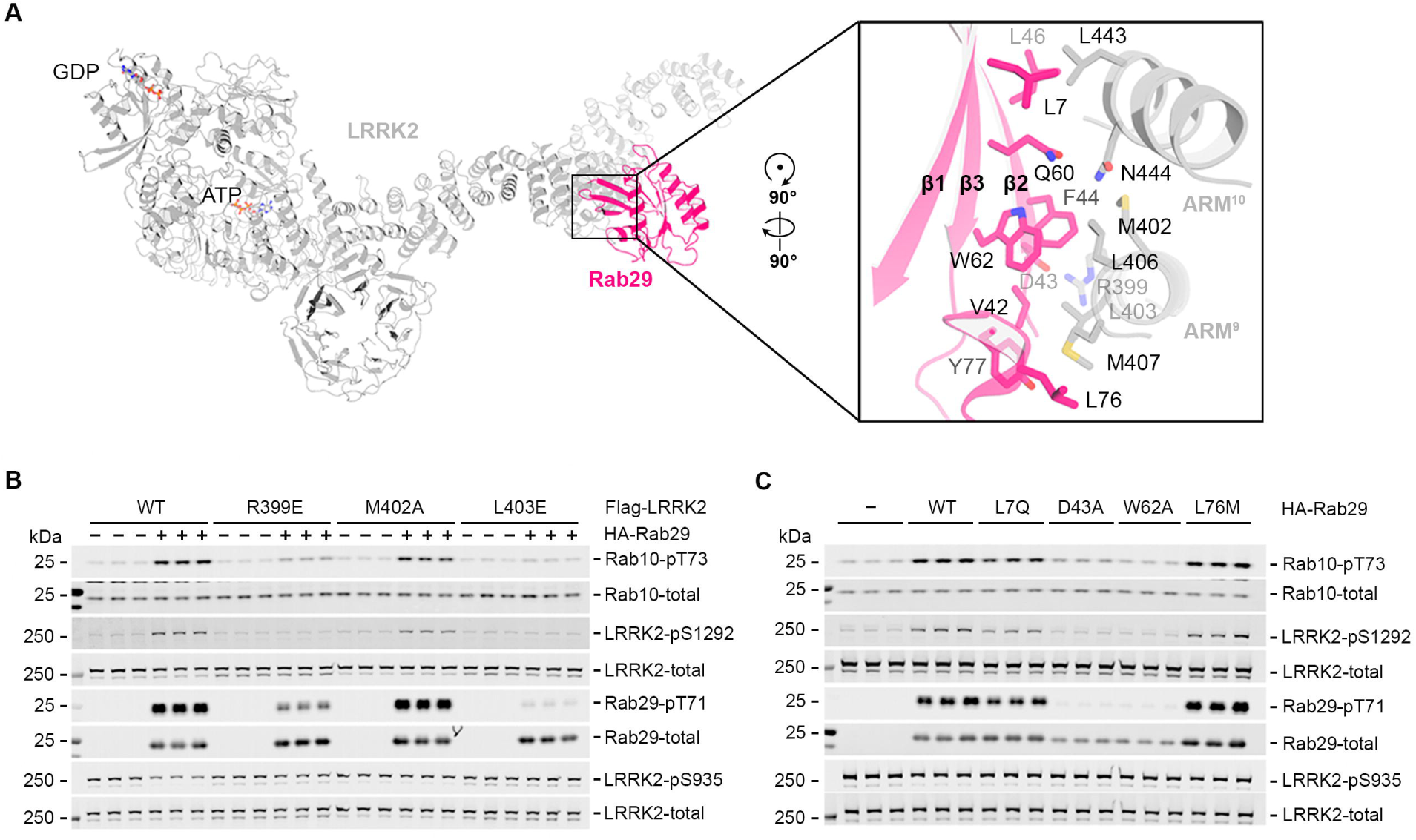
Structural and functional analysis of the LRRK2–Rab29 complex. **(A)** Structure of the LRRK2–Rab29 complex in the LRRK2 monomer state. LRRK2 and Rab29 are colored in grey and hot pink, respectively. Sidechains of the interface residues are shown as sticks. **(B)** Quantitative immunoblotting analysis of the cellular kinase activity of LRRK2 bearing mutations in the LRRK2/Rab29 interface. HEK293 cells were co-transfected with wild - type LRRK2 or the indicated variants and either HA-empty vector (−) or HA-tagged Rab29 (+). **(C)** Quantitative immunoblotting analysis of the cellular kinase activity of wild-type LRRK2 in the presence of Rab29 bearing mutations in the LRRK2/Rab29 interface. HEK293 cells were co-transfected with wild-type LRRK2 and either HA-empty vector (−) or the indicated variants of HA- tagged Rab29.

The LRRK2–Rab29 interface is formed between the ARM9-10 of LRRK2 and the “Switch I-Interswitch-Switch II” surface and complementarity determining region 1 (CDR1) (Pylypenko et al., 2018) of Rab29 (Figures S3A-B). This interaction conforms with a general Rab-effector recognition mode: effectors preferentially associate with the GTP-bound form of Rabs through the “Switch I-Interswitch-Switch II” surface (Pylypenko et al., 2018). Sequence alignment of Rab GTPase members, including Rab29, Rab32, Rab28 and several other LRRK2 substrates, reveals that key residues in the LRRK2–Rab29 interface are mostly conserved (Figure S3F). Moreover, introducing single mutations within the middle of the interface (Rab29^D43A^ or Rab29^W62A^) is sufficient to abolish the interaction between Rab29 and LRRK2 in GST pull-down assays. In contrast, Rab29^L7Q^ or Rab29^L76M^ mutations, localized at the edge of the binding interface, have minimal or no impact on the LRRK2–Rab29 association (Figure S3E). Intriguingly, Rab5A/B/C, which are LRRK2 kinase substrates under *in-vitro* overexpression conditions (Steger et al., 2017), contain an alanine in the corresponding position of Asp43 in Rab29 (Figure S3F), suggesting LRRK2 substrates might bind to a different site on LRRK2.

The effect of LRRK2–Rab29 interface mutations on LRRK2 kinase activity was examined in cells by monitoring the phosphorylation of Rab10 at Thr73, Rab29 at Thr71 and LRRK2 autophosphorylation at Ser1292 (Purlyte et al., 2018) (Figures 1B-C and S3G-H). Consistent with the GST pull-down assay, Rab29^D43A^ or Rab29^W62A^ mutations almost completely abolished the increased LRRK2 kinase activity induced by Rab29 overexpression, whereas Rab29^L7Q^ or Rab29^L76M^ had little or no effect (Figures 1C and S3H). Additionally, LRRK2 mutations at the LRRK2–Rab29 interface (LRRK2^R399E^, LRRK2^M402A^ or LRRK2^L403E^) also significantly reduced the kinase activities (Figures 1B and S3G).

### An inactive LRRK2 dimer state

In the LRRK2 dimer state structure, each protomer of LRRK2 binds a single Rab29 molecule at the ARM9-10 interface, as described above (Figure S1B). In this “X-shaped” complex, LRRK2 protomers adopt the same inactive conformation, as observed in the LRRK2 monomer state and the LRRK2 alone structure with an RMSD of 0.7 Å (Figures S4C-D) (Myasnikov et al., 2021). Notably, the interaction between two LRRK2 molecules in the LRRK2 dimer state is mediated by COR-B domains, similarly to what we described for the Rab29-free LRRK2 homodimer (Myasnikov et al., 2021). The observation of LRRK2 dimers both in the presence and absence of Rab29 leads us to believe the COR-B-mediated dimerization could occur under physiological settings, and the LRRK2 dimer state might serve as a transition from the inactive LRRK2 monomer state to the active LRRK2 tetramer state, described below.

### A tetrameric assembly of LRRK2

We determined an unexpected structure of the LRRK2 tetramer state to an overall resolution of 3.5 Å with C2 symmetry imposed (Figures S1-S2 and Table S1). With a ∼205 Å × 260 Å × 150 Å dimension, the LRRK2 tetramer state is an assembly with a two-fold rather than four-fold symmetry, featuring two peripheral and two central LRRK2 protomers (termed LRRK2^per^ and LRRK2^cen^ hereafter) (Figure 2). We resolved and modeled near full-length LRRK2^per^ and LRRK2^per^-associated Rab29 molecules (Rab29^per^). LRRK2^cen^ protomers have flexible LRR, ANK and ARM domains. Those domains, together with their possibly associated Rab29 (Rab29^cen^) molecules, could not be resolved due to flexibility (Figures 2 and S4A). On the other hand, the catalytic halves of LRRK2^cen^, including ROC, COR, KIN and WD40 domains, were rigid and refined to an overall resolution of 3.3 Å (Figure S1A).

**Figure 2.**
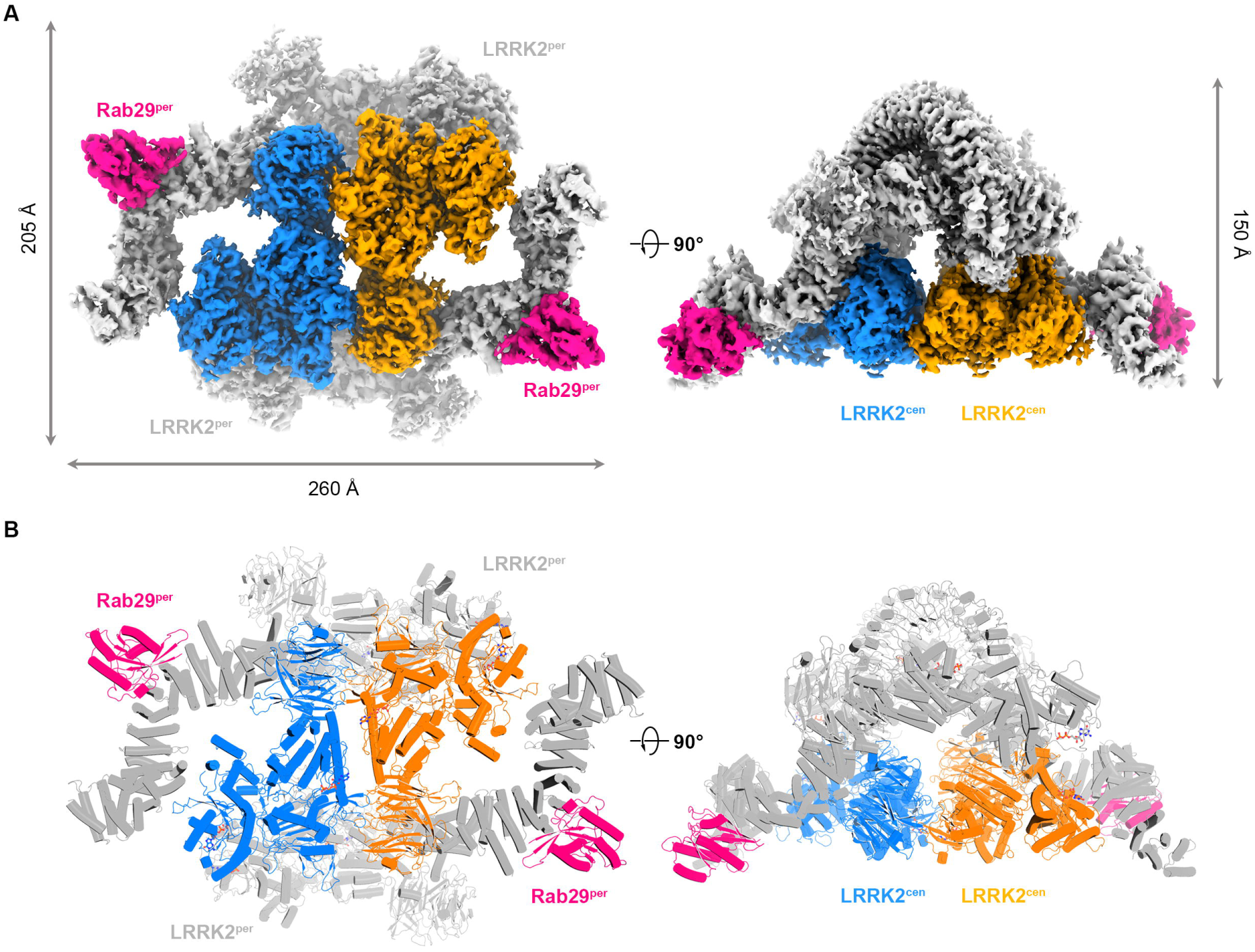
Cryo-EM structure of the LRRK2–Rab29 in the LRRK2 tetramer state. Cryo-EM map. **(A)** and model **(B)** of the LRRK2–Rab29 complex in the LRRK2 tetramer state with two different views. Peripheral Rab29 (Rab29^per^) and LRRK2 (LRRK2^per^) are colored in hot pink and grey, respectively; Central LRRK2 (LRRK2^cen^) copies are colored in blue and orange.

The LRRK2^per^–LRRK2^cen^ interaction within each asymmetric unit is mediated by the COR-B domains (Figures S4E-F), and the COR-B–COR-B interface is similar to that seen in LRRK2 homodimers (Myasnikov et al., 2021) or the LRRK2 dimer state (Figure S4F). LRRK2^per^ directly interacts with LRRK2^cen^ from the other asymmetric unit (Figure S4G), with the ARM domain of LRRK2^per^ associating with the ROC domain of LRRK2^cen^ through potential salt bridges, and regions near LRRK2^per^ ARM–ANK boundary packing against the WD40 domain of LRRK2^cen^ via hydrophobic interactions and hydrogen bonds (Figure S4G). The two LRRK2^cen^ protomers pack in a “head-to-tail” mode through WD40–KIN interfaces, and the C-terminal helices of the two LRRK2^cen^ protomers are antiparallel to each other (Figure S4G). The flexible N-terminal part of LRRK2^cen^ might also interact with the Rab29^per^ and the ARM domain of LRRK2^per^, but the low resolution of the cryo-EM map prevents further interpretation (Figure S4A).

### LRRK2 activation within the LRRK2 tetramer state

The catalytic halves of LRRK2^cen^ and LRRK2^per^ show substantial conformational differences. Upon aligning the COR-B domains, the KIN and WD40 domains are displaced about 40° towards the COR-A and ROC domains in LRRK2^cen^. This conformational rearrangement closes the central cavity shaped by ROC, COR and KIN domains (Figures 3A and S5B). Here ROC-COR-A, KIN^N-lobe^ and KIN^C-lobe^-WD40 appear to move as rigid bodies (Figure S5A). Repositioning of KIN^C-lobe^-WD40 in LRRK2^cen^ disrupts the connection between WD40 and ARM-ANK-LRR domains in the inactive state, which were bridged by the scaffolding “hinge helix” and “C-terminal helix” (Figures S6A-B) (Myasnikov et al., 2021) The KIN^C-lobe^ would also clash into the LRR domain (Figure S6C), which contributes to displacement and flexibility of the ARM-ANK-LRR domains within the LRRK2^cen^ protomers.

**Figure 3.**
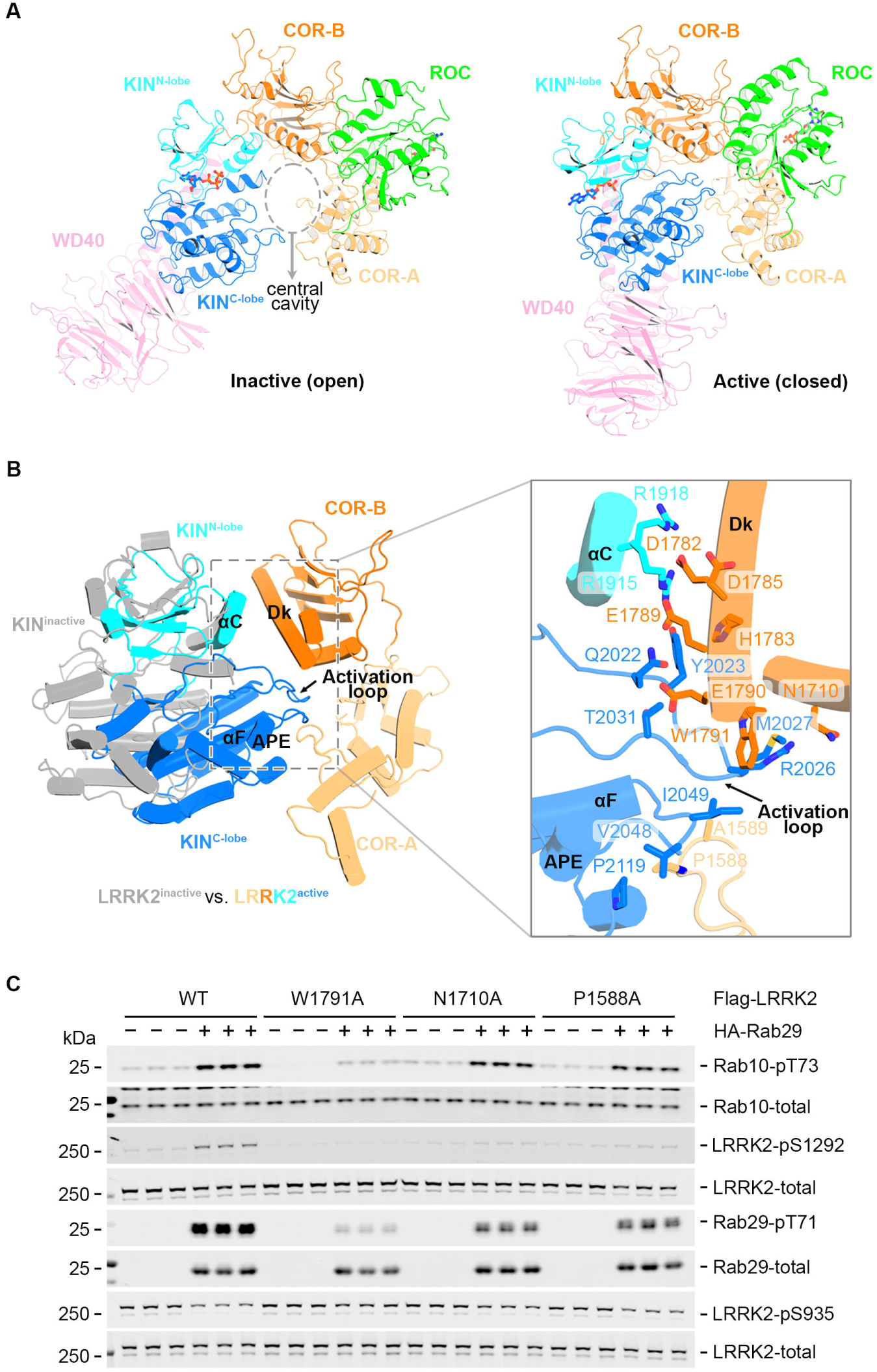
Conformational changes of LRRK2 upon activation. **(A)** Conformational changes in the C-terminal halves of LRRK2 upon activation. A dashed circle indicates the “central cavity” between the KIN and COR domains. Color codes for different parts of LRRK2 are as follows: ROC in green, COR-A in light orange, COR-B in bright orange, N-lobe of KIN in cyan, C-lobe of KIN in marine and WD40 in pink. The same color codes are used unless otherwise noted. **(B)** Movement of the KIN domain relative to the COR domain upon activation. Interactions between the KIN and COR domains in the active conformation are illustrated. Sidechains of the interface residues are shown as sticks. **(C)** Quantitative immunoblotting analysis of the cellular kinase activity of LRRK2 bearing mutations in the interface between the KIN and the COR domains. HEK293 cells were co-transfected with wild-type LRRK2 or the indicated LRRK2 variants and either HA-empty vector (−) or HA-tagged Rab29 (+).

In the LRRK2 tetramer state, the KIN domain of LRRK2^cen^ becomes exposed, manifesting features of an active kinase. The LRR domain that shields the KIN domain in the inactive conformation (Myasnikov et al., 2021) also becomes flexible (Figure S4A), leaving the KIN domain accessible to substrates. Critically, the LRRK2^cen^ KIN domain adopts a closed conformation (Figure 4A), with the αC helix positioned towards the active site and the “DYG-motif” flipped in. Lys1906 and Glu1920 form a salt bridge, an interaction blocked by Tyr2018 in the inactive conformation (Figures 4B-C). There is a well-defined cryo-EM density for ATP in the active site (Figure 4A), and the distance between Asp2017 and ATP shortens to 3.8 Å from 13.8 Å (Figures 4B-C), primed to permit ATP hydrolysis. Also, the regulatory spine (R-spine), formed by Leu1935, Leu1924, Tyr2018 and Tyr1992, becomes continuous (Figure 4D). These structural features point to the LRRK2^cen^ KIN domain being captured in an active conformation. The conclusion is further supported by the perfect docking of the LRRK2^cen^ model into the 14-Å *in-situ* cryo-ET map of LRRK2 (Figure S5F), which was reported to represent an active state (Deniston et al., 2020; Watanabe et al., 2020).

**Figure 4.**
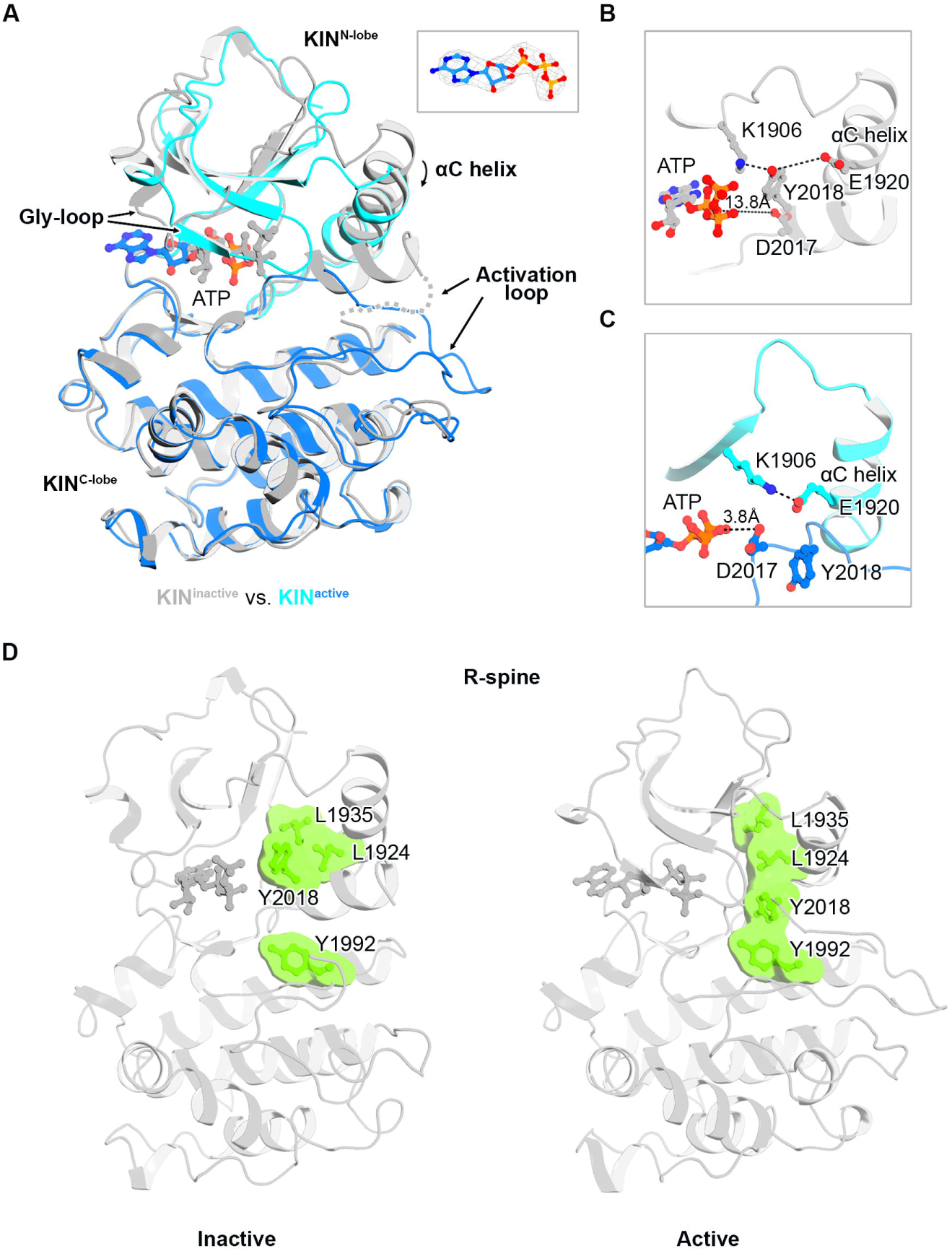
The active KIN domain of LRRK2^cen^. **(A)** Superposition of the active and inactive kinase domains. N- and C-lobes of the active kinase domain are colored in cyan and marine, respectively, and the inactive conformation is colored in grey. The cryo-EM density of the ATP molecule is shown. **(B)** Key catalytic residues in inactive kinase domain with sidechains shown as ball-and-stick models. The distance between the side chain of D2017 and the phosphate group of ATP is indicated with a dashed line. **(C)** Key catalytic residues in the active kinase domain. Interactions between K1906, E1920, D2017 and ATP are indicated by dashed lines. **(D)** The R-spine of the inactive (left) and active (right) KIN domains. The four residues forming the R-spine (L1935, L1924, Y2018, and Y1992) are shown as green surfaces.

Inter-domain interactions stabilize the active conformation of the LRRK2^cen^ KIN domain. The activation loop of the KIN domain dips into and enlarges the pocket between COR-A and COR-B (Figures 3B and S5E). We hypothesize that this interdomain interaction stabilizes the closed conformation of the KIN domain and is, thus, crucial for LRRK2 kinase activity. Indeed, point mutations of residues at the KIN–COR interface (P1588A, N1710A and W1791A) impact the increased kinase activity induced by Rab29 (Figures 3C and S5G). Therefore, we believe that blocking this interaction between COR and KIN domains could serve as a potential strategy to inhibit LRRK2 allosterically.

KIN^N-lobe^ rotates slightly towards the COR-B domain to form more intensive interactions between the αC helix of the KIN domain and the Dk helix of the COR-B domain in the active LRRK2^cen^ (Figures 3B and S5C). These structural observations are consistent with the previous hydrogen-deuterium exchange mass spectrometry (HDX-MS) and molecular dynamics (MD) simulation studies that indicated an altered interface between the COR-B Dk helix and KIN αC helix upon binding of type I inhibitors (Weng et al., 2022). Rotation of the KIN^N-lobe^ also stabilizes a COR-B loop (residues 1721–1725) that was disordered and exhibited a higher HDX rate in the inactive state (Figure S5C) (Weng et al., 2022).

The ROC domain is displaced relative to the COR-B domain upon LRRK2 activation (Figures S5B and S5D). COR-B structurally bridges the catalytic ROC and KIN domains, and the GTPase activity of the former modulates the kinase activity of the latter (Biosa et al., 2012; Ito et al., 2007; Taymans et al., 2011; West et al., 2007). Here we show that the movement of ROC relative to the COR-B domain upon activation has a striking “seesaw-like” motion of ROC’s αC helix, with Tyr1699 from COR-B (a PD-causing mutation site) being the pivot point (Figures 5D and S5D). Our structural observations support the proposal that conformational coupling between ROC and COR-B domains is important for LRRK2 kinase activity by contributing to the crosstalk between GTPase and kinase activities (Myasnikov et al., 2021).

**Figure 5.**
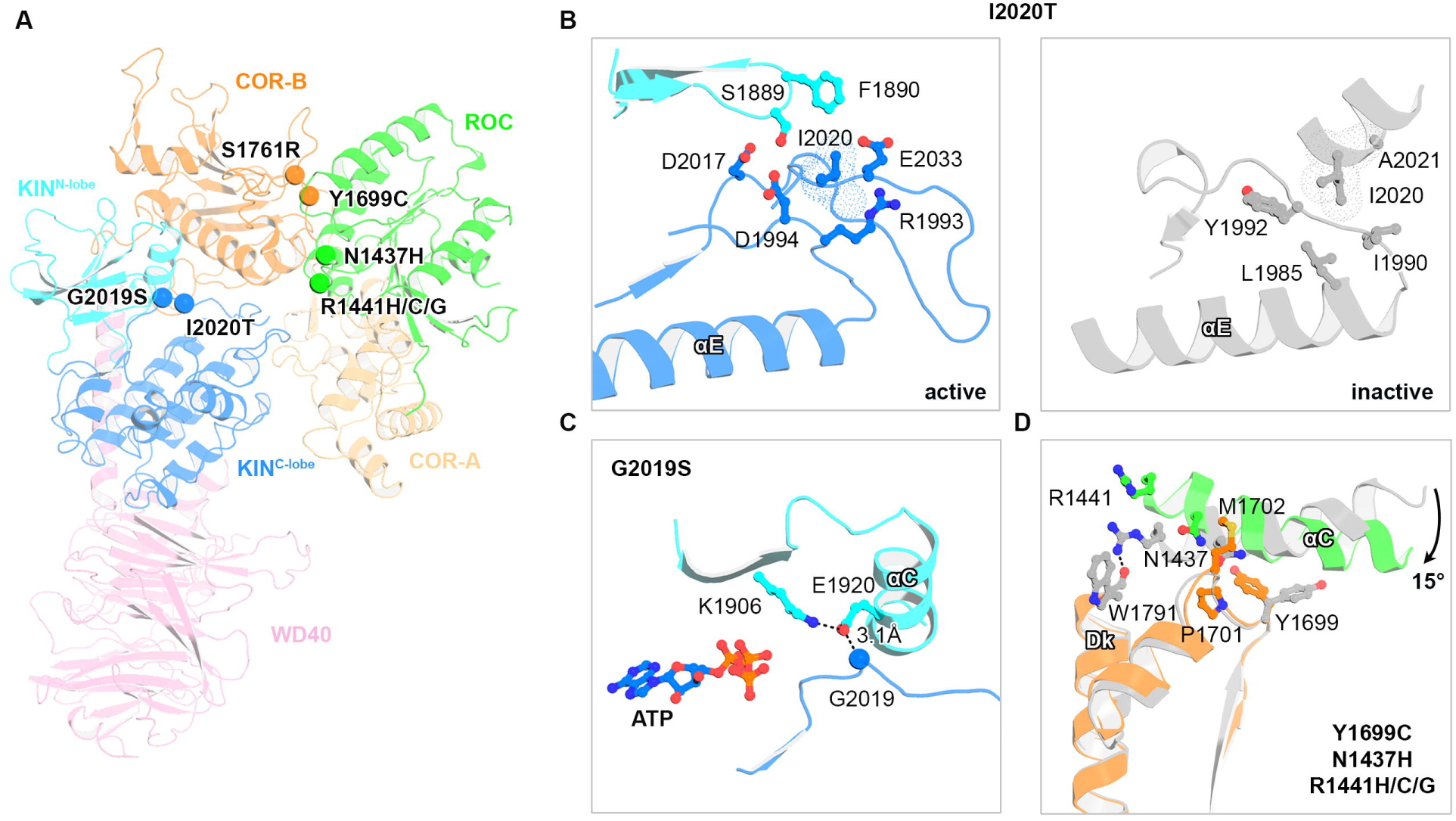
Structural mapping of disease-causing mutations. **(A)** Locations of PD-causing mutation sites in the active state of LRRK2. The mutation sites are shown as spheres and labeled. **(B)** The chemical environment of Ile2020 in active and inactive LRRK2. **(C)** Position of the Gly2019 residue relative to the K1906-E1920 salt bridge in the active LRRK2. The residue Gly2019 is shown as a sphere, and the distance between Gly2019 and Glu1920 is labeled. **(D)** Tyr1699, Asn1437, and Arg1441 sites at the ROC-COR interface. The motion of the αC helix in the ROC domain between inactive and active states of LRRK2 is indicated by a black arrow.

Our structural analysis reveals the molecular basis underlying the LRRK2 spatial regulation and kinase activation via the formation of a two-fold symmetric LRRK2 tetramer assembly. In this super-assembly, two LRRK2 protomers (LRRK2^cen^) become active, while the other two (LRRK2^per^) remain inactive. The LRRK2 tetrameric assembly, stabilized by both intra- and inter-subunit interactions, leads to massive conformational changes in the LRRK2^cen^ protomers, which results in flexible N-terminal ARM-ANK-LRR domains in the absence of membrane support and makes the KIN domain accessible to globular substrates (Figures S4A and 6A). More importantly, membrane anchoring of the lipidated Rab29^per^ molecules would orient active sites of exposed LRRK2^cen^ KIN domains towards the membrane to phosphorylate membrane-tethered proteins like prenylated Rab GTPases (Figure 6A), further supporting the physiological significance of the tetrameric assembly of LRRK2.

**Figure 6.**
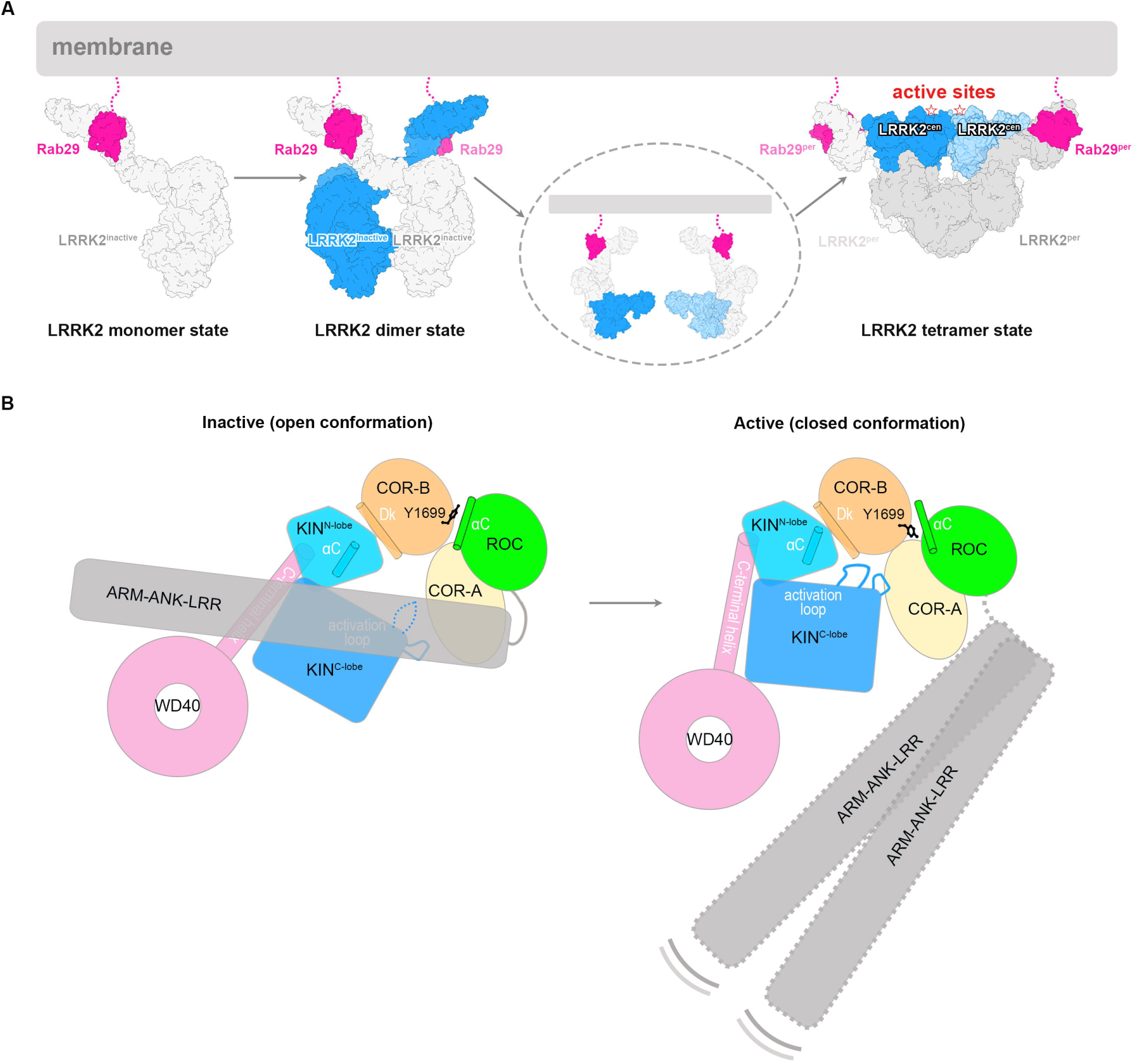
Proposed model of LRRK2 activation by Rab29. **(A)** Working model for Rab29-mediated LRRK2 activation. Rab29 is colored in hot pink, and the lipid anchor is indicated by dashed lines. LRRK2 is colored grey in the LRRK2 monomer state and grey and blue in the LRRK2 dimer state. In the LRRK2 tetramer state, two LRRK2^per^ are colored in grey, and two LRRK2^cen^ are colored in blue. The concentration-dependent oligomerization and activation processes are indicated by arrows. **(B)** A cartoon showing major conformational differences between inactive and active LRRK2.

### Structural interpretation of LRRK2 PD mutations

Having captured the LRRK2 in both active and inactive states now allows us to analyze PD mutations in the context of LRRK2 activation. Gain-of-function mutations at six sites—G2019S, I2020T, Y1699C, N1437H, R1441C/G/H and S1761R (Figure 5A)—have been proposed to be high-risk and PD-causing (Christensen et al., 2018). Substitution of Gly2019 with serine showed little structural difference from the wild type in the inactive state (Myasnikov et al., 2021) but is expected to induce additional interactions with Glu1920 and stabilize the critical Lys1906-Glu1920 salt bridge in the active kinase conformation (Figure 5C). Ile2020 moves from a hydrophobic to a hydrophilic environment upon LRRK2 activation (Figure 5B), so the LRRK2^I2020T^ mutation destabilizes the inactive conformation and favors the active conformation. Tyr1699, Asn1437 and Arg1441 are found at the interface between the αC helix of ROC and the COR-B domain (Figure 5D), where a “seesaw-like” motion of ROC’s αC helix occurs upon LRRK2 activation (Figures 5D and S5D). Asn1437 and Arg1441 are located at one side of the seesaw and anchor the C-terminal part on the αC helix to the surface of the COR-B domain in the inactive state (Figure 5D). Therefore, mutations of Asn1437 and Arg1441 would weaken the anchoring effect, shift the balance of the “seesaw” towards the N-terminal of the ROC αC helix and promote kinase activation. Tyr1699 of the COR-B domain, as noted above, directly functions as the pivot point for the “seesaw” (Figure 5D). Thus, we propose that substituting Tyr with a smaller residue would lower the energy barrier for the “seesaw” motion and the transition from the inactive to the active state.

## Discussion

Here we report LRRK2–Rab29 complex structures in the LRRK2 monomer, dimer and tetramer states; with structure-guided biochemical analyses, our work presents a novel kinase activation paradigm, where LRRK2 activation is controlled by Rab29-dependent spatial recruitment and nonsymmetric tetramerization of LRRK2. Recent studies have suggested an emerging “Rab29-LRRK2-Rab8A/Rab10” cascade (Kuwahara and Iwatsubo, 2020). Specifically, lipidated Rab29 recruits LRRK2 to the *trans*-Golgi network (TGN), where LRRK2 oligomerizes and becomes enzymatically active, resulting in the phosphorylation of Rab substrates, such as Rab8A and Rab10. Our study provides structural snapshots with atomic details illustrating possible key steps during this activation cascade, summarized in Figure 6A. In this model, prenylated Rab29 recruits LRRK2 via its ARM domain, and membrane recruitment leads to increased local concentration and COR-B-mediated LRRK2 dimerization. Then, Rab29-bound dimeric LRRK2 could undergo Rab29-induce, oligomerization-associated conformational changes to form the LRRK2 tetramer state with a dimer-of-asymmetric dimer assembly (Figure 6A). In the LRRK2 tetramer state, two of the four LRRK2 protomers are kinase-active and expose active sites towards the membrane to phosphorylate membrane-attached substrates like lipidated Rab8A and Rab10 (Figure 6A). Moreover, the LRRK2 tetrameric assembly fits perfectly with the expected membrane geometry and previous biochemical data that imply the importance of LRRK2 oligomerization for its kinase activity (Berger et al., 2010; Civiero et al., 2017; Gloeckner et al., 2006; Greggio et al., 2008; James et al., 2012; Jorgensen et al., 2009; Sen et al., 2009).

The Rab29- and oligomerization-dependent activation of LRRK2 reveals a novel kinase activation mechanism. LRRK2 activates through a striking dimer-of-asymmetric dimer assembly containing two active core subunits encased by two inactive peripheral protomers. Intensive intermolecular interactions between two asymmetric LRRK2 dimers stabilize the active LRRK2 tetramer state (Figure S4G). We believe that the formation of active LRRK2 tetramers is Rab29-dependent since such an assembly was not observed in the absence of Rab29 under otherwise similar experimental conditions (Myasnikov et al., 2021). The binding of Rab29 to the extended ARM-ANK-LRR portion of LRRK2^cen^ appears to unlock or facilitate the conformational changes that lead to the active LRRK2^cen^ conformation (Figure S4A), but the low resolution of LRRK2^cen^ ARM-ANK-LRR domains prevents us from dissecting such possibilities in detail. Nevertheless, our work reveals a dual role of Rab29 in both LRRK2 recruitment and activation. This Rab29- and oligomerization-controlled kinase activation mechanism is novel and unprecedented to the best of our knowledge. It is distinct from the well-known asymmetric kinase activation mechanism, where kinase domains form asymmetric dimers to become functionally active, such as EGFR (Zhang et al., 2006) and B-Raf (Kondo et al., 2019). Here we show a novel paradigm of non-symmetric kinase activation mechanism, in which the Rab GTPase signal and protein oligomerization synergistically modulate the functionality of a protein kinase.

The flexible ARM-ANK-LRR domains of LRRK2^cen^ may contribute to substrate recruitment. LRRK2 specifically recognizes and phosphorylates a subset of Rab GTPases (Steger et al., 2017; Steger et al., 2016). Our biochemical data suggest the presence of a potential secondary Rab binding site (Figure S3F). If this site is located in ARM-ANK-LRR domains, we postulate that these domains can potentially use the flexibility to recruit substrates.

In addition to revealing new structural insights into known pathogenic PD mutations, our work also highlights new opportunities for the rational design of allosteric LRRK2 inhibitors. For example, our data indicate that allosteric inhibition of LRRK2 could be achieved by disrupting LRRK2–Rab29 interaction, by blocking LRRK2 oligomerization or by preventing the conformation transition from inactive to active states. These findings provide a solid knowledge foundation for the rational design of allosteric inhibitors of LRRK2. Altogether, we have captured the LRRK2-Rab29 complex in three different conformational states and provide key structural snapshots that likely occur during LRRK2 membrane recruitment and activation. The Rab29-dependent activation of LRRK2, which reveals an unexpected mechanism of coupling membrane recruitment with protein oligomerization, unveils a novel kinase activation paradigm. Thus, our work sets the stage for the interpretation of disease-causing mutations in the context of LRRK2 activation and paves the way for allosteric inhibitor design for PD treatment.

## Acknowledgments

We thank Yu-Hua Lo of the Cryo-electron Microscopy Center at St. Jude Children’s Research Hospital for help with cryo-EM data collection, members of the Sun lab for helpful discussions, Norman Luo for bio-illustration and Ines Chen, Jian Payandeh for valuable inputs for manuscript preparation. We also thank the excellent technical support of the MRC Protein Phosphorylation and Ubiquitylation Unit (PPU) DNA cloning team (coordinated by Rachel Toth), sequencing service (coordinated by Gary Hunter), and tissue culture team (coordinated by Edwin Allen). This work was funded by the American Lebanese Syrian Associated Charities (ALSAC), NIH (R00HL143037), the UK Medical Research Council (MC_UU_00018/1) and Aligning Science Across Parkinson’s (ASAP). The Michael J. Fox Foundation for Parkinson’s Research (MJFF) administers the grant (ASAP-000463) on behalf of ASAP.

## Author Contributions

HZ and JS designed the project. HZ performed sample preparation, biochemical analysis and structural determination. FT designed, performed and analyzed the cellular activity assays. FT and DRA interpreted the activity assays. All authors were involved in data analysis. HZ and JS wrote the manuscript with input from all authors.

## Figure legends

**Figure S1.** Structure determination of the LRRK2–Rab29 complexes.

**(A)** A simplified flow chart of the cryo-EM data processing.

**(B)** The overall structure of LRRK2-Rab29 complexes in three different oligomerization states.

**Figure S2.** Data and model quality assessment for the LRRK2–Rab29 complexes dataset.

**(A), (D) and (G)** Local resolution of LRRK2–Rab29 complexes in the LRRK2 monomer, dimer and tetramer states.

**(B), (E) and (H)** Fourier Shell Correlation (FSC) curves for the overall resolution of the LRRK2–Rab29 complexes in the LRRK2 monomer, dimer and tetramer states. The golden standard (FSC=0.143) is used for resolution estimation.

**(C), (F) and (I)** Model-to-map fit between the full map and PDB coordinates of the LRRK2–Rab29 complexes in the LRRK2 monomer, dimer and tetramer states.

**Figure S3.** Structural features of the LRRK2–Rab29 complex in the LRRK2 monomer state

**(A)** Cartoon model of Rab29 with secondary structures and key structural elements labeled. The phosphorylation sites are shown as spheres.

**(B)** The interaction between the LRRK2 ARM domain and Rab29. Key secondary structures involved in protein-protein interactions are labeled.

**(C)** Superposition of LRRK2-bound Rab29 with other Rab proteins in GTP- and GDP-bound states. “Switch I” motifs are indicated by black lines.

**(D)** Superposition of Rab29 in the LRRK2–Rab29 complex with GDP-bound Rab29 (PDB code 6HH2). The steric clash between the “Switch I” motif of GDP-bound Rab29 and LRRK2 is indicated.

**(E)** *In-vitro* GST pull-down assays with wild-type (Q67L/T71A/S72A) or mutant GST-Rab29 and wild-type LRRK2. Gels were stained using Coomassie Blue.

**(F)** Sequence alignment of Rab29 subfamily members (Rab29, Rab32, and Rab38) and Rab substrates of LRRK2. The residues involved in the interaction with LRRK2 are indicated with cyan circles. The secondary structures and “switch” motifs are also labeled.

**(G)** and **(H)** Quantification of the immunoblotting data shown in Figures 1B and 1C, respectively. Data are presented as ratios of pRab10-Thr73/total Rab10, pLRRK2-Ser1292/total LRRK2, and pRab29-Thr71/total Rab29, normalized to the average of LRRK2 wild-type values (mean ± SD).

**Figure S4.** Structural features of the LRRK2–Rab29 complex in the LRRK2 tetramer state.

**(A)** Density map for the LRRK2–Rab29 complex in a low threshold showing potential density of the N-terminal part of LRRK2^cen^ molecules.

**(B), (C)** and **(D)** Structural comparison of the C-terminal catalytic halves of LRRK2 between inactive LRRK2 (PDB 7LHW) (Myasnikov et al., 2021) and LRRK2^per^ in the LRRK2 tetramer state, LRRK2 in the LRRK2 monomer and dimer states, respectively.

**(E)** Structural comparison of the LRRK2–Rab29 of the LRRK2 dimer state and asymmetric unit of the LRRK2 tetramer state in two different views.

**(F)** Superposition of COR-B mediated LRRK2 dimer interfaces between the LRRK2 dimer and tetramer states. The conformational change is indicated by an arrow.

**(G)** Interactions between LRRK2^cen^ and LRRK2^per^ copies. **Left**: Interactions between the ROC domain of LRRK2^cen^ and ARM domain of LRRK2^per^. **Right**: Interactions between the WD40 domain of LRRK2^cen^ and ARM/ANK domain of LRRK2^per^.

**Figure S5.** The active closed conformation of LRRK2.

**(A)** Structural comparison of the rigid bodies in the C-terminal catalytic halves between inactive and active LRRK2.

**(B)** Superposition of the C-terminal catalytic halves of the inactive and active LRRK2 copies. The conformational changes are indicated with arrowed lines.

**(C)** Comparison of COR-B subdomain and KIN^N-lobe^ between the inactive and active LRRK2 copies. The rotational movement of the KIN^N-lobe^ towards COR-B and the loop (1721-1725) undergoing conformational changes are indicated. The color code is the same as Figure S5B.

**(D)** Comparison of ROC-COR-B domains between the inactive and active LRRK2 copies. The rotational movement of the ROC domain relative to the COR-B subdomain is indicated. The proposed “seesaw-like” model constituted by Tyr1699 of COR-B subdomain and αC helix of ROC domain is shown in a zoom-in window.

**(E)** Comparison of COR-A and COR-B subdomains between the inactive and active LRRK2 copies. The rotational movement of the COR-A subdomain relative to the COR-B subdomain is indicated.

**(F)** Docking of the C-terminal catalytic half of active LRRK2 into the cryo-ET map of microtubule-bound LRRK2^I2020T^ (Watanabe et al., 2020).

**(G)** Quantification of the immunoblotting data shown in Figure 3C (Validation of the COR/KIN interface by cell-based kinase activity assays). Data are presented as ratios of pRab10-Thr73/total Rab10, pLRRK2-Ser1292/total LRRK2, and pRab29-Thr71/total Rab29, normalized to the average of LRRK2 wild-type values (mean ± SD).

**Figure S6.** The flexibility of the N-terminal half of LRRK2 upon activation.

**(A)** Interdomain interactions in inactive LRRK2 (Myasnikov et al., 2021). **Left**: the key scaffold elements: “hinge helix” and “C-terminal helix” are indicated. **Right**: the three interaction sites between KIN and LRR domains are shown in dashed circles.

**(B)** The displacement of WD40 domain and C-terminal helix upon activation.

**(C)** The movement of the KIN domain upon activation. The potential steric clashes between the C-lobe of KIN and LRR domains upon activation are indicated by a dashed circle.

## Method Details

### Cloning, expression and purification of human LRRK2 and Rab29

The cDNA encoding human LRRK2 was purchased from Horizon Discovery and contains three natural variants (R50H, S1647T and M2397T) from the canonical LRRK2 sequence (NP_940980.4). A GFP tag followed by a preScission protease cleavage site was engineered at the N terminus of LRRK2, which was cloned into the BacMam expression vector (Goehring et al., 2014). Recombinant baculoviruses of LRRK2 were generated using the Bac-to-Bac system according to the manufacturer’s instructions (Invitrogen). Then P3 virus of LRRK2 was used for transfection of HEK293F cells for protein expression. Briefly, for 1L cultures of HEK293F cells (∼2-3×10^6^ cells/mL) in Freestyle 293 media (Gibco) supplemented with 2% FBS (Gibco), about 100 ml P3 virus was used. Infected cells were incubated at 37 °C overnight, and protein expression was induced by adding 10 mM sodium butyrate. Cells were cultured at 30 °C for another 48-60 hrs before harvest. The cell pellet from the 600 mL culture was resuspended in 30 ml lysis buffer (20 mM Tris pH 8.0, 200 mM NaCl, 10% glycerol, 2 mM DTT and protease inhibitors), and then cells were lysed by brief sonication. LRRK2 was separated from the insoluble fraction by high-speed centrifugation (38,000 g for 1 hr), and incubated with 1 ml CNBr-activated sepharose beads (GE Healthcare) coupled with 1 mg high-affinity GFP nanobodies (GFP-NB) (Kirchhofer et al., 2010). The GFP tag was cleaved by preScission protease at 4 °C, and LRRK2 was further purified by size-exclusion chromatography with a Superose 6 Increase 10/300 GL column (GE Healthcare) equilibrated with 20 mM Tris pH 8.0, 200 mM NaCl and 2 mM DTT. The purified protein was collected and concentrated to 13 mg/ml (OD_280_) using a 100-kDa MWCO centrifugal device (Ambion), flash-frozen in liquid N_2_ and stored at -80 °C.

The cDNA encoding human Rab29 was purchased from Horizon Discovery. Residues 1-177 of human Rab29 were cloned into the modified pGEX plasmid (Sigma), which attaches a GST tag and a TEV protease cleavage site at the N-terminus. For structural and biochemical studies, we generated Rab29^Q67L/T71A/S72A^ mutant, which was constructed using the QuickChange Site-Directed Mutagenesis kit (Strategene). The recombinant proteins were overexpressed in *Escherichia coli* strain BL21 (DE3) in LB media supplemented with 0.05 mg/ml ampicillin, and cells were grown at 37 °C until OD_600_ reached 0.8. Further expression was induced by adding 0.4 mM IPTG, and cells were allowed to grow at 16 °C for 20 hrs. Harvested cells were lysed by sonication in the lysis buffer (20 mM Tris-HCl pH 8.0, 200 mM NaCl, 5% glycerol, 5 mM MgCl_2_, and 1 mM PMSF), and lysates were cleared by centrifugation at 38,000 g for 45 min. Subsequently, the proteins were purified by GST affinity chromatography on the glutathione Sepharose beads (GE Healthcare) in lysis buffer. The bound proteins were then eluted with lysis buffer containing 20 mM GSH and incubated with 1 mM GTP at 4 °C overnight followed by gel filtration chromatography using a Superdex 200 Increase 10/300 GL column (GE Healthcare) in a storage buffer (20 mM Tris-HCl pH 8.0, 100 mM NaCl, 1 mM MgCl_2_, and 2 mM DTT). For structural study, the GST tag was removed by on-column cleavage with TEV protease at 4 °C overnight before gel filtration chromatography. The peak fractions were collected and concentrated to 16 mg/ml (OD_280_), flash-frozen in liquid N_2,_ and stored at -80 °C.

### CryoEM analysis

CryoEM grids were prepared with a Vitrobot Mark IV (FEI). Quantifoil R1.2/1.3 300 Au holey carbon grids (Quantifoil) were glow-discharged for 30 s. Purified LRRK2 and Rab29 were incubated together on ice for 1 hr with a final concentration of 30 μM and 200 μM, respectively. In addition, 2.3 mM fluorinated Fos-Choline-8 were added right before freezing the grids and then 3.0 μL of protein sample was pipetted onto the grids, which were blotted for 5 s under blot force - 3 at 95% humidity and frozen in liquid nitrogen-cooled liquid ethane.

CryoEM data were collected using Titan Krios (Thermo Fisher) transmission electron microscope, equipped with a K3 direct electron detector and post-column GIF (energy filter). K3 gain references were collected just before data collection. Data collection was performed in an automatic manner using EPU software (Thermo Fisher Scientific). Movies were recorded at defocus values from -0.6 to -1.8 μm at a magnification of 81kx, which corresponds to the pixel size of 1.06 Å at the specimen level (super-resolution pixel size is 0.53). During 3.0-second exposure, 60 frames (0.05 s per frame and the dose of 0.9795 e/frame/Å^2^) were collected with a total dose of ∼59 electrons Å^-2^. In total, 11,762 images were collected. Motion correction was performed on raw super-resolution movie stacks and binned by 2 using MotionCor2 software (Zheng et al., 2017). Contrast transfer function (CTF) estimation was performed using Gctf (Zhang, 2016). Prior particles picking micrographs were analyzed for good power spectrum, and the bad ones were discarded (with 11,260 good images remaining).

Particles were selected using the template picker in cryoSPARC (Punjani et al., 2017) and extracted using a binning factor of 2. Several rounds of the 2D classification were performed to eliminate ice, carbon edges, and false-positive particles containing noise. During 2D classification, three groups of classes were observed, corresponding to the LRRK2 monomer, dimer and tetramer states, respectively. All groups were selected, and *ab initio* reconstruction was performed. In order to further separate LRRK2 monomer, dimer and tetramer states, we perform Heterogeneous Refinement in cryoSPARC. As a result, 104,718 particles were assigned to the monomer class, 86,379 particles to the dimer, and 62,946 particles to the tetramer. All 3D classes were refined in parallel using cryoSPARC after extraction of corresponding particles without binning. For monomer, we performed a standard NU-refinement by applying C1 symmetry followed by a focused 3D refinement to improve the map quality of the ARM domain of LRRK2 with Rab29 binding. As to dimer, we performed an NU-refinement by applying C2 symmetry. For tetramer, in addition to an NU-refinement by applying C2 symmetry, we also perform a symmetry expansion procedure in order to get a better resolution for the peripheral part and a focused refinement for the central part in order to further improve resolution. Local and global CTF refinements were performed to improve the resolution of the final reconstruction. All resolution was reported according to the gold-standard Fourier shell correlation (FSC) using the 0.143 criterion (Henderson et al., 2012). Local resolution was estimated in cryoSPARC. The density maps sharpened in cryoSPARC were used to produce figures.

### Model building and refinement

The reported structures of LRRK2 (PDB codes: 7LI4 and 7LHT) (Myasnikov et al., 2021) and the predicted structures of LRRK2 and Rab29 by AlphaFold2 (Jumper et al., 2021) were fitted and adjusted into the cryo-EM maps of LRRK2 monomer, dimer, and tetramer using Chimera (Pettersen et al., 2004) and Coot (Emsley et al., 2010). The structural model was refined against the map by using the real space refinement module with secondary structure and non-crystallographic symmetry restraints in the Phenix package (Adams et al., 2010). Fourier shell correlation curves were calculated between the refined model and the full map. The geometries of the models were validated using MolProbity (Chen et al., 2010). All the figures were prepared in PyMol (Schrödinger, LLC.), UCSF Chimera (Pettersen et al., 2004) and UCSF ChimeraX (Goddard et al., 2018).

### GST pull-down assay

Twenty micrograms (or 0.4 nmol) of WT or various mutants of GST-tagged Rab29 proteins were immobilized onto 30 μl of the glutathione Sepharose beads (GE Healthcare) and then incubated with 870 μg (or 3 nmol) LRRK2 proteins in 1 ml binding buffer containing 20 mM Tris-HCl pH 8.0, 100 mM NaCl, 10% glycerol, 1 mM GTP, 5 mM MgCl_2_, 2 mM DTT and protease inhibitors for 3 hrs at 4 °C. The beads were washed three times (10 min each) with binding buffer, and bound proteins were eluted with binding buffer supplemented with 20 mM GSH. The protein samples were subjected to SDS-PAGE followed by Coomassie brilliant blue staining. GST was used as the control.

### Plasmids used for cell-based kinase activity assays

The following plasmids were used for the cell-based kinase activity assays: HA-empty vector (DU49303); HA-Rab29 wild-type (DU50222); HA-Rab29 L7Q (DU72178); HA-Rab29 D43A (DU72179); HA-Rab29 W62A (DU 72177); HA-Rab29 L76M (DU 72180); Flag-LRRK2 wild-type (DU62804); Flag-LRRK2 R399E (DU72192); Flag-LRRK2 M402A (DU72193); Flag-LRRK2 L403E (DU72194); Flag-LRRK2 W1791A (DU72200); Flag-LRRK2 N1710A (DU72212); Flag-LRRK2 P1588A (DU72201). All these plasmids were generated by the MRC PPU Reagents and Services at the University of Dundee (https://mrcppureagents.dundee.ac.uk). Each LRRK2 variant was confirmed by sequencing at the MRC Sequencing and Services (https://www.dnaseq.co.uk). All plasmids are available to request via the MRC PPU Reagents and Services website (https://mrcppureagents.dundee.ac.uk).

### Cell transfection and lysis

A detailed description of our cell transfection method (dx.doi.org/10.17504/protocols.io.bw4bpgsn) and cell lysis method (dx.doi.org/10.17504/protocols.io.b5jhq4j) has previously been described. Briefly, HEK293 cells were seeded into 6-well plates and transiently transfected at 60-70% confluence using Polyethylenimine (PEI) transfection reagent (Polysciences, Inc., #24765). For each well, 1.6 μg of N-ter Flag-tagged LRRK2 (wild-type or mutant), 0.4 μg of N-ter HA-tagged Rab29 (wild-type or mutant) (or HA-empty vector) and 6 μg of PEI were diluted in 0.5 mL of Opti-MEM™ Reduced serum medium (Gibco™) and incubated for 15-20 minutes at room temperature before being added to the cells. Cells were lysed 24 hrs post-transfection in ice-cold lysis buffer containing 50 mM Tris-HCl pH 7.4, 1 mM EGTA, 10 mM 2-glycerophosphate, 50 mM sodium fluoride, 5 mM sodium pyrophosphate, 270 mM sucrose, supplemented with 1 μg/ml microcystin-LR, 1 mM sodium orthovanadate, complete EDTA-free protease inhibitor cocktail (Roche), and 1% (v/v) Triton X-100. Lysates were clarified by centrifugation at 17,000 g at 4 °C for 10 min and supernatants were quantified by Bradford assay.

### Quantitative immunoblot analysis

A detailed description of our quantitative immunoblotting protocol has previously been described (dx.doi.org/10.17504/potocols.io.bsgrnby6). Briefly, cell lysates were mixed with a quarter of a volume of 4 x SDS-PAGE loading buffer (Invitrogen™ NuPAGE™ LDS Sample Buffer, cat# NP0007) and heated at 70 °C for 5 min. Samples were loaded onto NuPAGE 4–12% Bis–Tris Midi Gels (Thermo Fisher Scientific, Cat# WG1402BOX or Cat# WG1403BOX) and electrophoresed at 130 V for 2 hrs in NuPAGE MOPS SDS running buffer (Thermo Fisher Scientific, Cat# NP0001-02). Proteins were then electrophoretically transferred onto a nitrocellulose membrane (GE Healthcare, Amersham Protran Supported 0.45 μm NC) at 90 V for 100 min on ice in transfer buffer (48 mM Tris base and 39 mM glycine supplemented with 20% (v/v) methanol). The membranes were blocked with 5% (w/v) skim milk powder dissolved in TBS-T (50 mM Tris base, 150 mM sodium chloride (NaCl), 0.1% (v/v) Tween 20) at room temperature for 1 hr before overnight incubation at 4 °C in primary antibodies. Membranes were washed three times for 15 min each with TBS-T before being incubated with secondary antibodies for 1 hr at room temperature. Thereafter, membranes were washed with TBS-T three times with a 15-minute incubation for each wash, and protein bands were acquired via near-infrared fluorescent detection using the LI-COR Odyssey CLx Western Blot imaging system.

### Antibodies used for quantitative immunoblotting analysis

The antibodies against Rab10 pThr73 [MJF-R21] (ab230261), LRRK2 pSer1292 [MJFR-19-7-8] (ab203181) and Rab29 pThr71 [MJF-R24-17-1] (ab241062) were purchased from Abcam. The rabbit monoclonal antibody against LRRK2 pSer935 (UDD2) was purified by MRC PPU Reagents and Services at the University of Dundee (https://mrcppureagents.dundee.ac.uk). The mouse monoclonal antibody against total LRRK2 (C-terminus) was purchased from NeuroMab (clone N241A/34, #75-253). The mouse monoclonal against total Rab10 was purchased from Nanotools (#0680–100/Rab10-605B11). The anti-HA tag antibody was purchased from Sigma (cat #11867423001). All primary antibodies were diluted in 5% (w/v) bovine serum albumin (BSA) in TBS-T and used at a final concentration of 1 μg/ml.

Goat anti-mouse IRDye 680LT (#926-68020), goat anti-rabbit IRDye 800CW (#926-32211) and goat anti-rat IRDye 680LT (#926-68029) IgG (H + L) secondary antibodies were from LI-COR and were diluted 1:20,000 (v/v) in 5% (w/v) milk in TBS-T.

## Quantification and Statistical Analysis

Immunoblotting data for the cell-based kinase activity assays shown in Figures 1B-C and 3C were quantified using Image Studio software. Quantified data were plotted with GraphPad Prism.The values are averages of three independent determinations with standard deviations. The quantification and statistical analyses for model refinement and validation were generated using MolProbity (Chen et al., 2010).

