## Supplementary figures and images for "Structural basis of human LRRK2 membrane recruitment and activation"

### Figure S1

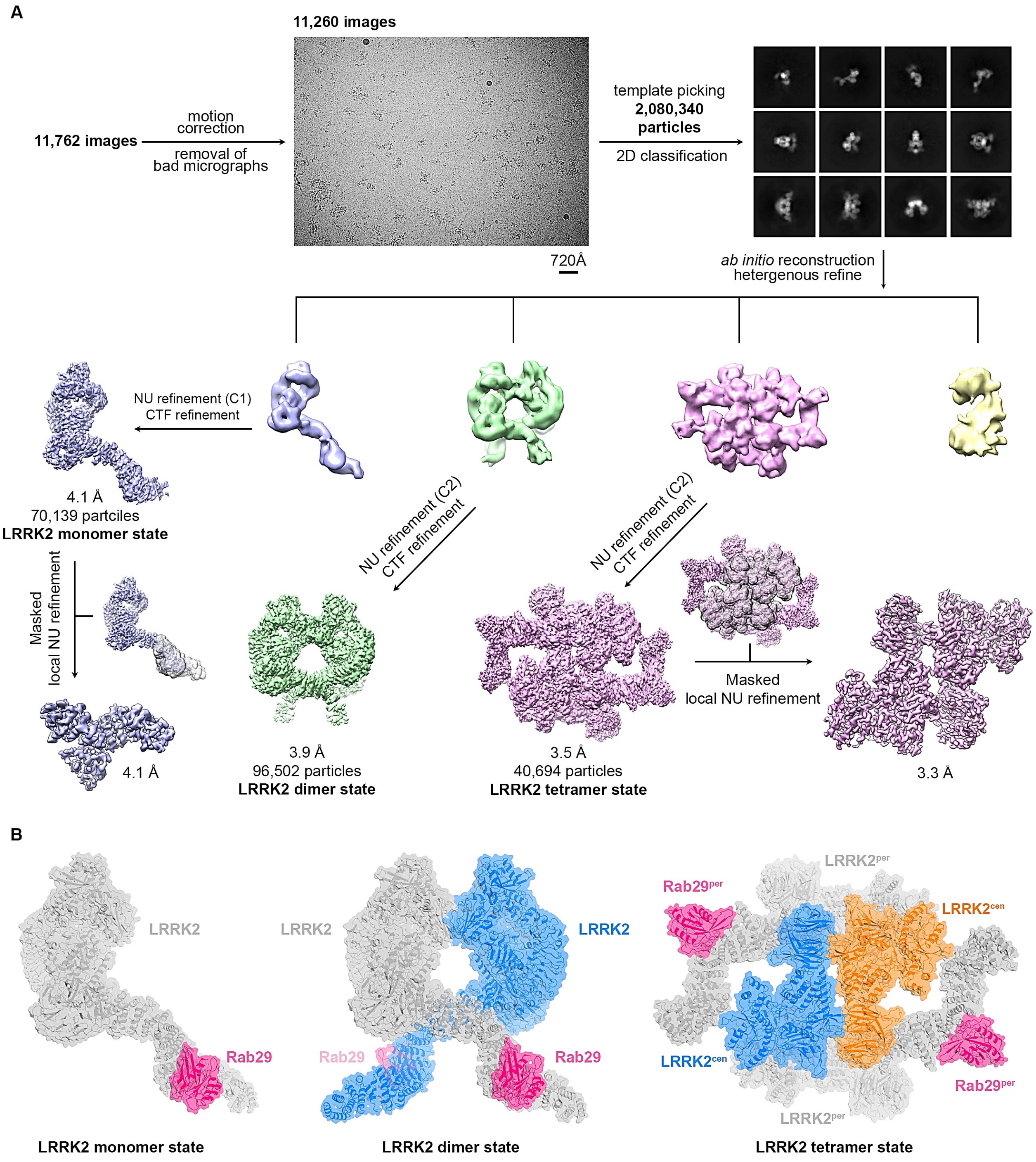

### Figure S2

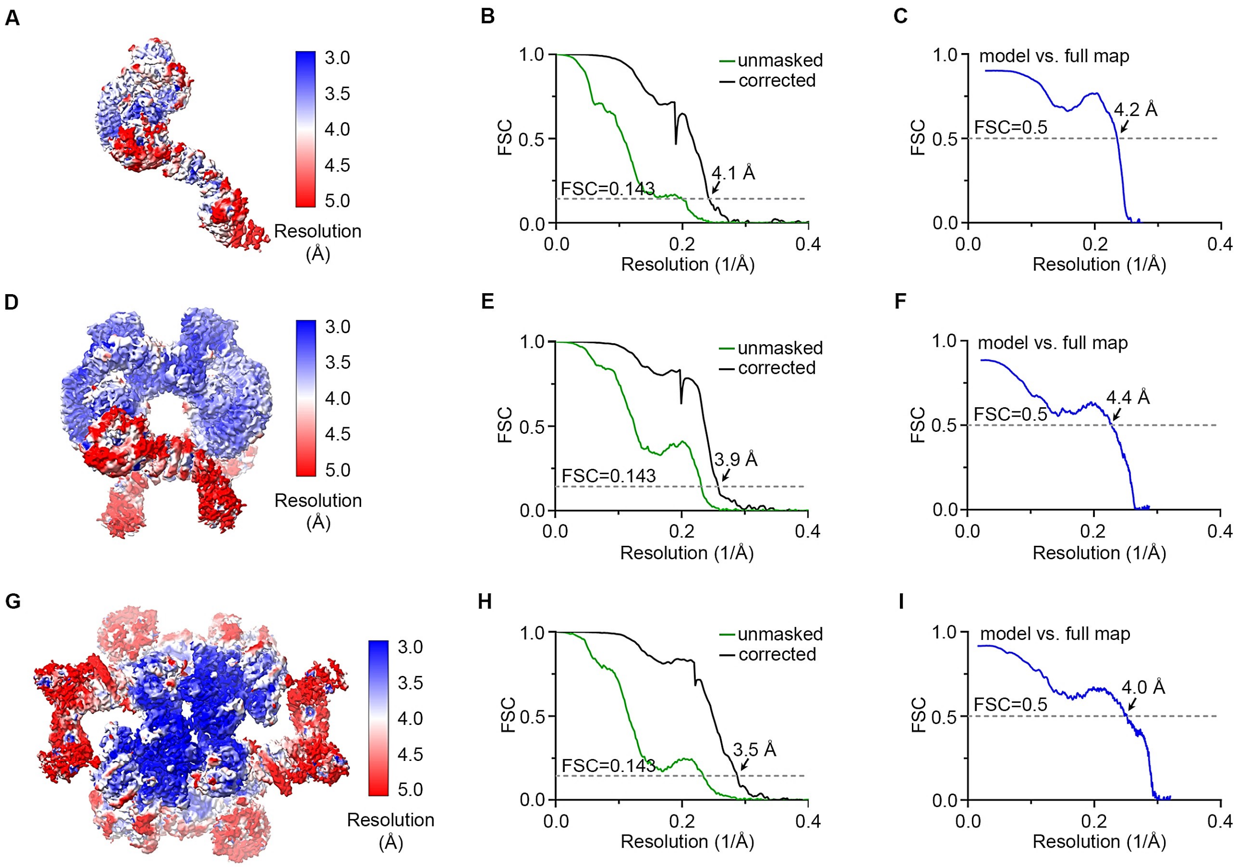

### Figure S3

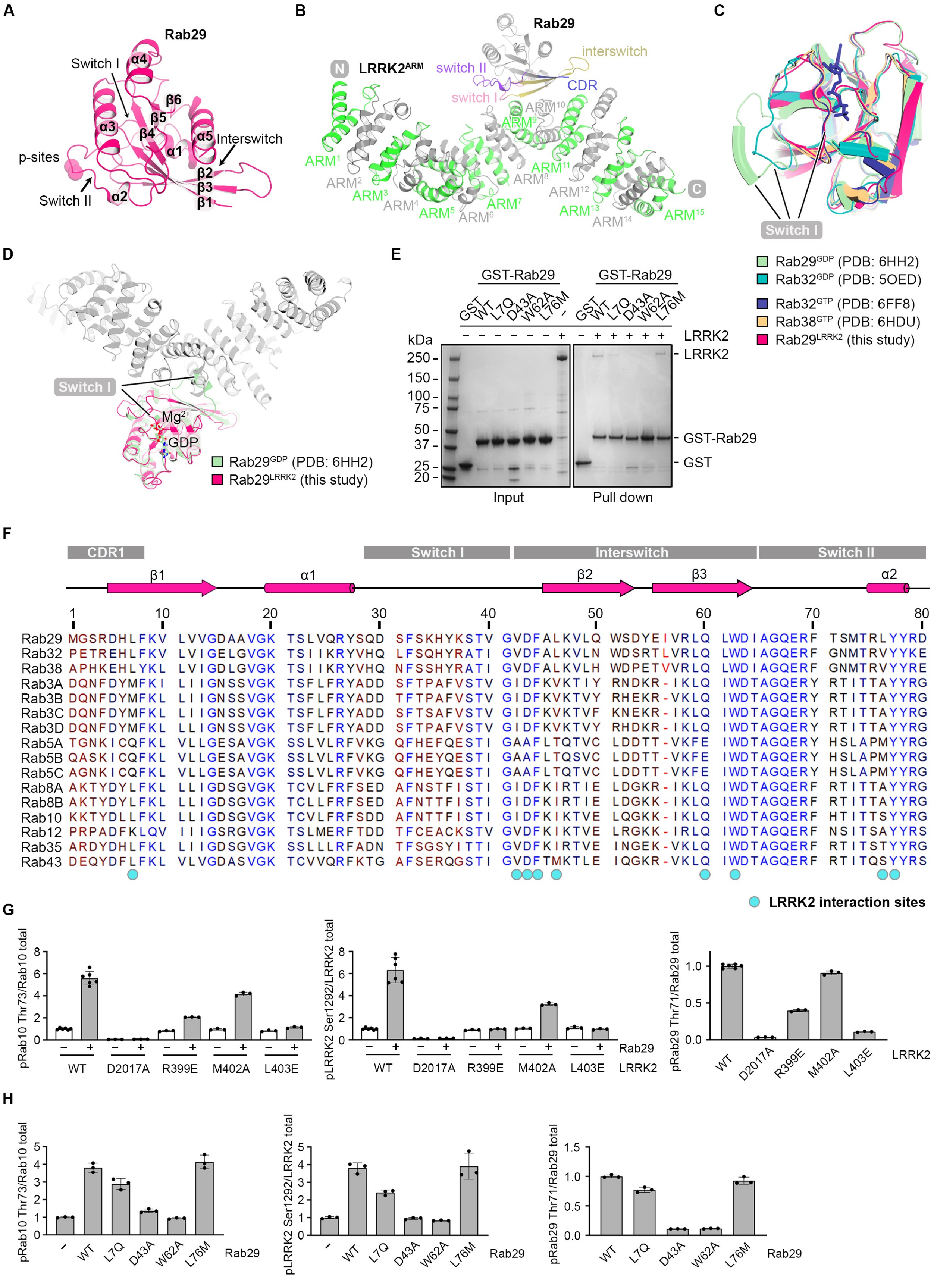

### Figure S4

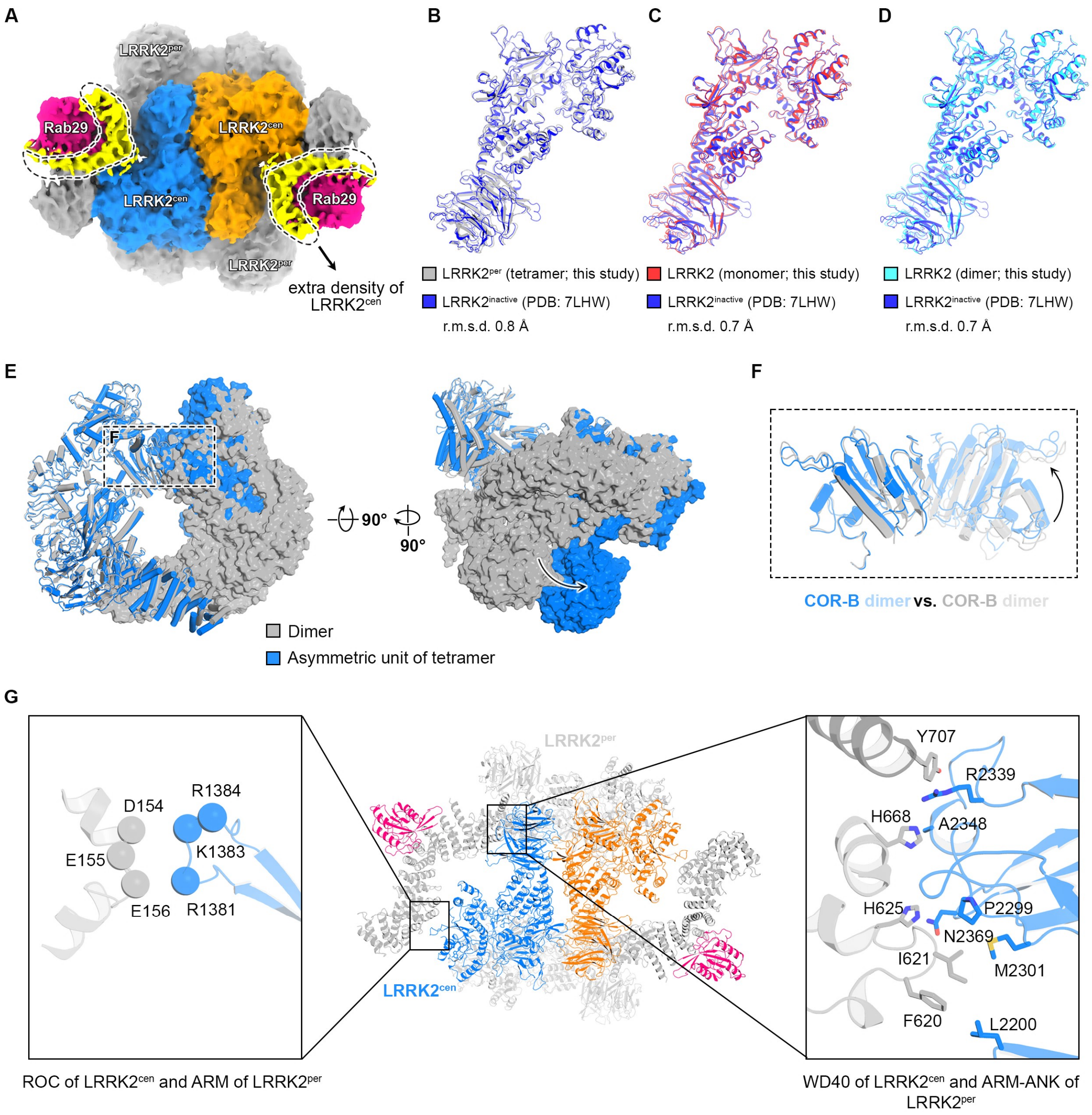

### Figure S5

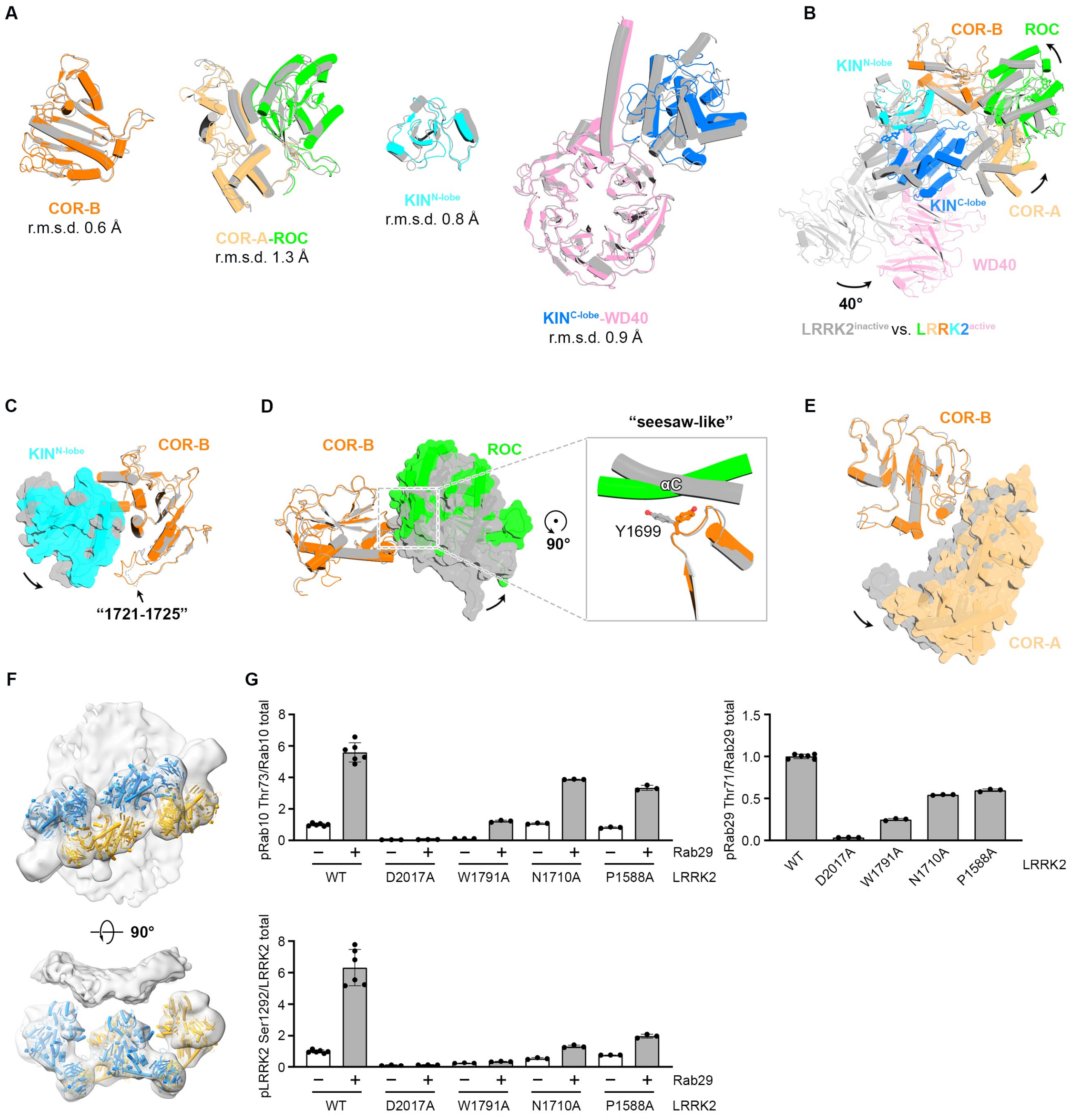

### Figure S6

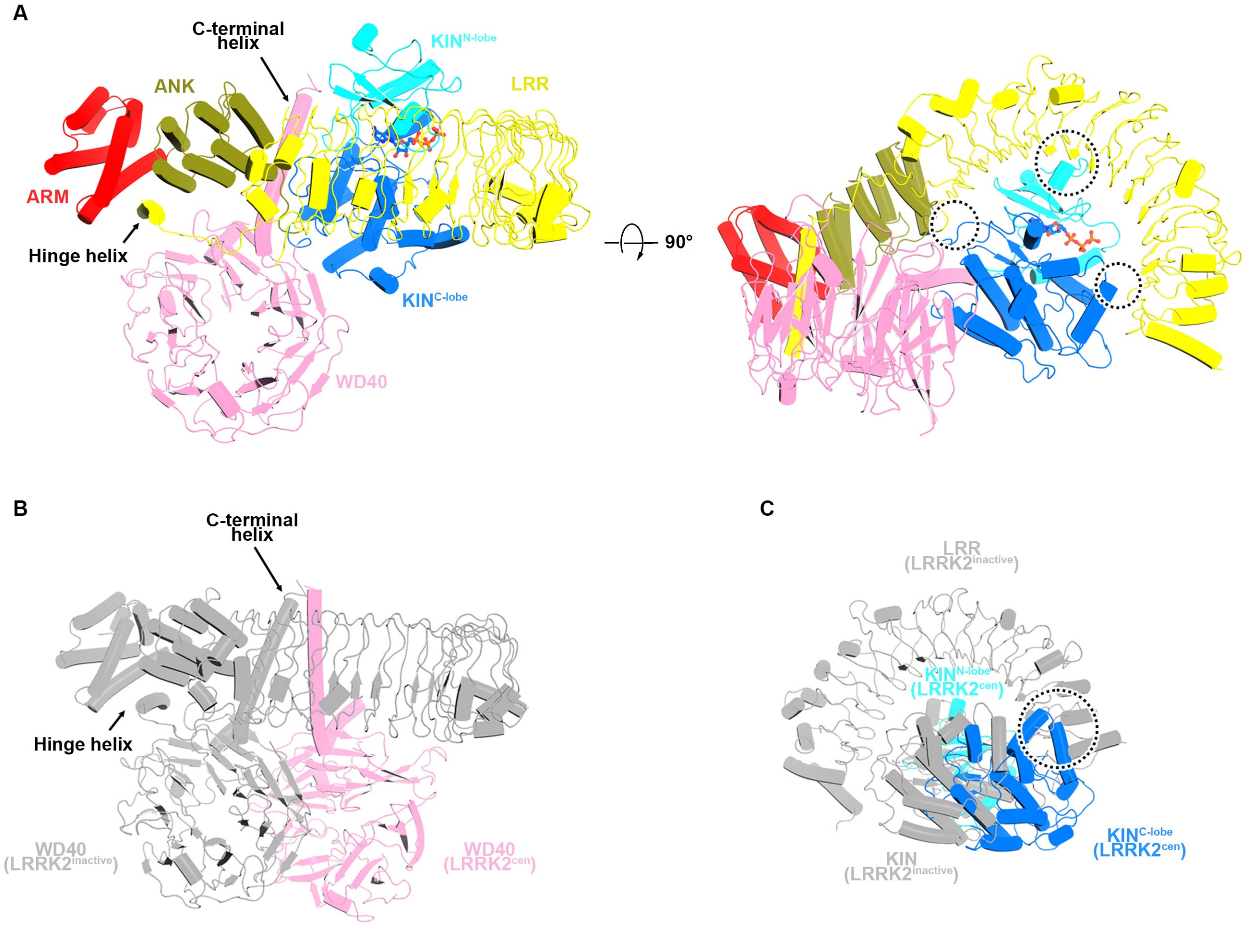
