## Supplementary material for "Structural basis of human LRRK2 membrane recruitment and activation": Table S1

Table S1. Cryo-EM data collection, refinement and validation statistics

| | LRRK2-Rab29 monomer PDB XXXX EMD-XXXXX | LRRK2-Rab29 dimer PDB XXXX EMD-XXXXX | LRRK2-Rab29 tetramer PDB XXXX EMD-XXXXX |
|---|---|---|---|
| **Data collection and processing** | | | |
| Microscope/Camera | | Titan krios/Gatan K3 Camera | |
| Magnification | 81,000 | 81,000 | 81,000 |
| Voltage (kV) | 300 | 300 | 300 |
| Electron exposure ($e^-/Å^2$) | 58.77 | 58.77 | 58.77 |
| Defocus range (μm) | 0.6-1.8 | 0.6-1.8 | 0.6-1.8 |
| Pixel size (Å) | 1.06 | 1.06 | 1.06 |
| Symmetry imposed | C1 | C2 | C2 |
| Initial particle images (no.) | 354,085 | 249,148 | 186,624 |
| Final particle images (no.) | 70,139 | 96,502 | 40,694 |
| Map resolution (Å) | 4.13 | 3.88 | 3.48 |
| FSC threshold | 0.143 | 0.143 | 0.143 |
| **Refinement** | | | |
| Initial model used (PDB code) | 7LHW and 6HH2 | XXXX | XXXX |
| Model resolution (Å) | 4.3 | 4.2 | 3.8 |
| FSC threshold | 0.5 | 0.5 | 0.5 |
| Map sharpening *B* factor ($Å^2$) | -135.2 | -120.0 | -82.5 |
| Model composition | | | |
| Non-hydrogen atoms | 18,231 | 36,452 | 54,304 |
| Protein residue atoms | 18,172 | 36,334 | 54,068 |
| Ligand atoms | 59 | 118 | 236 |
| *B* factors ($Å^2$) | | | |
| Protein | 130.8 | 130.8 | 89.0 |
| Ligand | 92.7 | 92.7 | 68.9 |
| R.m.s deviations | | | |
| Bond lengths (Å) | 0.003 | 0.004 | 0.003 |
| Bond angles (°) | 0.655 | 0.706 | 0.619 |
| **Validation** | | | |
| MolProbity score | 2.04 | 2.08 | 2.01 |
| Clashscore | 11.67 | 13.04 | 10.78 |
| Poor rotamers (%) | 0.00 | 0.00 | 0.07 |
| Ramachandran plot | | | |
| Favored (%) | 92.83 | 92.81 | 92.77 |
| Allowed (%) | 7.17 | 7.19 | 7.23 |
| Disallowed (%) | 0.00 | 0.00 | 0.00 |
